# Macrophage-derived WNT regulates tumor immune microenvironment to reduce colitis-associated colon cancer

**DOI:** 10.1101/2025.09.27.679002

**Authors:** Payel Bhanja, Rishi Man Chugh, Shujah Rehman, Stacey Krepel, Amrita Mitra, Pooja Gupta, Ximena Diaz Olea, Harsh Pathak, Anup Kasi, Subhrajit Saha

**Affiliations:** Departments of Radiation Oncology, University of Kansas Medical Center, Kansas City, KS, 66160, USA; Department of Pathology and Laboratory Medicine, University of Kansas Medical Center, Kansas City, KS 66160, USA; Department of Medicine Clinical Oncology, University of Kansas Medical Center, Kansas City, KS 66160, USA; Department of Cancer Biology, University of Kansas Medical Center, Kansas City, KS 66160, USA

**Keywords:** Colitis associated colon cancer, Macrophages, GSK3β, WNT, Tumor, PD1, PDL1

## Abstract

Prolonged colonic inflammation and ulcerative colitis lead to colon cancer. The rapid growth and treatment-resistant nature of these tumors are primarily influenced by an immunosuppressive tumor microenvironment, which is led by tumor-associated macrophages (TAMs). However, factors influencing or regulating the immunosuppressive nature of TAMs have not been sufficiently studied. In this manuscript, we use a mouse model of colitis-associated colorectal cancer (CRC) to demonstrate that WNT expression in TAMs regulates their immunosuppressive function by inhibiting Glycogen synthase kinase-3 beta (GSK-3β) within the macrophages, possibly through an autofeedback loop. GSK3β is a positive regulator of PD-1 and PDL1 expression in macrophages and promotes an immunosuppressive microenvironment. Therefore, GSK-3β inhibition alters the immunosuppressive nature of the immune microenvironment and effectively controls tumor growth. In Csf1r-iCre; Porcnfl/fl mice, the absence of macrophage-derived WNT promotes tumor growth in the model of colitis-associated colon cancer. Absence of macrophage-derived WNT stabilizes GSK-3β in macrophages and promotes an immunosuppressive tumor microenvironment. We also show that pharmacological inhibition of GSK-3β in a macrophage-specific manner, achieved by systemic delivery of a lipo-GSK3β inhibitor, effectively inhibits tumor growth. Therefore, this manuscript demonstrates for the first time that the macrophage-specific modulation of GSK3β can be a potential target to treat colitis-associated colon cancer.

## Introduction

Colorectal cancer (CRC) is one of the leading causes of death globally [1]. Persistent mucosal inflammatory disorder such as Crohn’s disease or Ulcerative colitis propelled the colorectal cancer risk. Patients with inflammatory bowel disease, for example ulcerative colitis and Crohn’s disease, are at an elevated risk for developing colon cancer [2, 3]. The molecular pathogenesis of colitis-associated cancer (CAC) remains incompletely understood. Inflammation and colorectal cancer are associated with increased inflammatory myeloid cell recruitment and function.

The immunosuppressive nature in the CRC tissue is not created solely by tumor cells but it is also mediated by non-tumoral stromal cells, most notably tumor associated macrophages (TAMs). These tumor-reprogramed macrophages are different from those in acute inflammatory responses where they phagocytose antigens, present antigens to and stimulate adaptive immune cells. Instead, TAMs shut down effector T cell activities with soluble immunosuppressive factors and membrane-bound immune checkpoint molecules such as the PD ligand 1 (PDL1) [4, 5]. Several studies demonstrated that tumor cell-derived regulatory factors that repolarize TAMs [6]. However, the mechanism by which TAMs integrate external signals and translate them into an immunosuppressive transcript has not been studied well. In this manuscript in mice model of colitis induced CRC we demonstrated that WNT expression in TAM regulates the immunosuppressive function of TAM by inhibiting GSK3beta within TAM possible by an auto feedback loop. Glycogen synthase kinase-3 β (GSK-3β) is a positive regulator of PD-1 expression in CD8+ T cells and GSK-3 inhibition enhances T cell function and is effective in the control of tumor growth [7]. In macrophages it has been shown that GSK3β-deficiency inhibits the progression of hepatocellular carcinoma and enhances the sensitivity of anti-PD1 immunotherapy [8]. Macrophage is one of the major sources of WNT ligand. Binding of WNT ligands with its receptor inhibits GSK-3 β. Numbers of publications over the years suggested that WNT signaling is critical oncogenic signal and very predominant in colon cancer. There have been several trials conducted to perturbed WNT/β-catenin signaling against colon cancer with moderated success. Our study in this manuscript demonstrated that activation of WNT signaling in macrophages is critical for regulation colitis induced colon cancer. *Csf1r-iCre; Porcn^fl/fl^* mice the absence of macrophage derived WNT promote tumor growth in mice model of colitis associated colon cancer with the stabilization of GSK-3β and promoting immunosuppressive tumor microenvironment. Pharmacological inhibition of GSK-3β in macrophage specific manner by systemic delivery of Lipo-GSK3β inhibitor inhibits the tumor growth.

## Materials and Methods

### Animals

To study the role of macrophage-derived WNT in colitis-associated colorectal cancer (CAC), *Porcn*, the gene encoding Porcupine, an O-acyltransferase essential for palmitoylation and secretion of WNT proteins, was deleted in macrophages by ablating a floxed allele of *Porcn* with Cre recombinase expressed under the colony-stimulating factor-1 receptor (Csf1r) promoter (Csf1r.iCre) [9]. *Csf1r.iCre; Porcnfl^/fl^* mice were generated by crossing floxed *Porcn* (*Porcn^fl/fl^*) mice with transgenic Csf1r.iCre mice, resulting in macrophage-restricted deletion of *Porcn* and loss of WNT secretion. Littermate *Porcn^fl/fl^* mice carrying intact floxed alleles were used as wild-type (WT) controls. Eight to ten weeks old female mice of both *Csf1r.iCre; Porcnfl^/fl^* and *Porcn^fl/fl^* genotypes were used in this study to develop the AOM/DSS model of colitis-associated colon cancer. All animals were maintained ad libitum, and experiments were conducted in accordance with protocols approved by the Institutional Animal Care and Use Committee (IACUC) of the University of Kansas Medical Center (ACUP#2020-2562).

### Development of a mouse model of colitis-associated colon cancer

Colon cancer development was initiated with a single intraperitoneal injection of azoxymethane (AOM; 6.5 mg/kg, body weight) (Sigma-Aldrich, Cat no. A5486). Starting 5 days after AOM administration, the mice received 3% DSS (36,000–50,000 MW., Colitis Grade, MP Biomedicals) in the drinking water for 7 days (Cycle-1), followed by a 2 week of without DSS water, then another 7 days cycle of 3% DSS (Cycle-2) followed by 2 weeks without DSS water and, subsequently, a final 7 day cycle of 2 % DSS (Cycle-3). This repeated DSS cycle develops a robust model of colitis-associated colorectal cancer with tumor development observed within 7–10 weeks. Mice were monitored daily and euthanized following the third DSS cycle for isolation of the colon to assess the tumor burden, immune checkpoint expression, and histological analysis. The colon tissues were also isolated at intermediate time points after each DSS cycle.

### DAI Scoring

Disease activity was assessed every other day using a Disease Activity Index (DAI) score derived from three parameters: body weight loss, stool consistency, and rectal bleeding. From Day 3 onwards, a Hemoccult test was performed on every alternate day to evaluate occult blood in the stool. This test was omitted if gross rectal bleeding or visibly red stool was observed.

The DAI score was calculated by summing the scores from the following criteria:

#### Body weight loss

0: No weight loss; 1: 1%-5% weight loss; 2: 5%-10% weight loss; 3: 10%-20% weight loss; 4: >20% weight loss

#### Stool consistency

0: Normal; 2: Loose stool; 4: Diarrhea

#### Rectal bleeding

0: No blood (Hemoccult negative); 2: Fecal occult blood test positive (Hemoccult positive); 4: Gross bleeding (visible blood around the anus)

### Colonoscopy and tumor count

A colonoscopy was performed to assess the tumor burden in the mouse colon using Karl Storz Endoskope, a specialized miniature endoscopic system for small animals. This consists of a miniature endoscope, a xenon light source for illumination, a high-definition camera, and an air pump to gently inflate the colon to avoid colon perforation and improve visibility during the imaging procedure. The endoscope camera transmits images of the bowel lining to the attached monitor, allowing for real-time visualization and detailed examination. Saline irrigation was used when necessary to clear any obscuring fecal matter or debris to improve the clarity of the endoscopic view. The endoscopic procedures enable repeated examinations of the same animals over time, providing data on disease progression and response to the treatments. It also helps in identifying the polyps and early cancerous lesions, facilitating early intervention and investigation of early-stage disease mechanisms.

Upon euthanasia, the colon was carefully taken out and opened longitudinally to assess the tumor burden. Tumors larger than 1 mm in diameter were counted on the colon’s luminal surface. The size of each tumor in a mouse is recorded, and the sum of all tumor sizes is calculated to obtain a tumor score, providing an overall measure of tumor burden. Half of the colon tissue was fixed in 10% neutral-buffered formalin for paraffin embedding and histological examination. The remaining half was processed for stromal cell isolation, flow cytometric analysis, and subsequent sorting for RNA sequencing and qPCR analysis.

### Histology

The colon of each animal was dissected, washed in PBS to remove luminal contents, opened longitudinally, and fixed in 10% neutral-buffered formalin before paraffin embedding. Tissue was routinely processed, and 5-μm sections were prepared for hematoxylin and eosin (H&E) and IHC staining. All H&E staining was performed at the Pathology Core Facility at the University of Kansas Cancer Center (Kansas City, KS).

For transmission electron microscopy (TEM), part of the fresh colon tissue was fixed with 2.5% glutaraldehyde. Tissue was washed 3 times with PBS at room temperature and post-fixed for 1 hour at 4°C in 1% osmium tetroxide with 1% potassium ferricyanide. Tissue samples were then dehydrated using a series of graded ethanol solutions, followed by a propylene oxide (PO) rinse and embedded in resin molds. Sectioning (80nM size) of the tissue was done using a diamond knife and Leica UC-7 and imaged in a JEOL JEM-1400 TEM at 100KV transmission electron microscope fitted with a Lab6 filament. **Immunohistochemistry**

For Ki67 and CD3 staining, colon tissue sections were deparaffinized and rehydrated with a graded series of alcohols, and colon sections were incubated overnight with a monoclonal anti-Ki67 antibody (M7240 mib1; Dako) and anti-CD3 antibody (14-0032-82, Invitrogen). Nuclear staining was detected using streptavidin-peroxidase with diaminobenzidine (DAB) as the chromogen, followed by counterstaining with hematoxylin. Quantification of Ki67 and CD3 positivity was performed by manually counting Ki67 and CD3-positive cells in at least ten randomly selected high-power fields per section, using an image captured with a Nikon NIS Element 80i microscope. The positive cells were expressed as a percentage of the total counted cells per field.

For immunofluorescence (IF) analysis, tissue sections were deparaffinized, and rehydrated colon sections were treated for antigen retrieval in citrate buffer (pH 6.0) at 95°C for 30 minutes. After cooling, sections are blocked with 5% normal goat serum in PBS for 1 hour at room temperature followed by incubation with Primary antibodies targeting DCLK (Thermo; Lab Vision), PD1 (Novus Biologicals), PDL1 (Invitrogen; MAS-20343), T-bet (Invitrogen # Cat # 14-5825-82), phospho-PTEN (Invitrogen# 44-1066G), phosphor-AKT(cell signalling; #9271), phosphor-GSK3beta (Invitrogen; # MAS14873 overnight at 4°C. Control sections without primary antibody to evaluate non-specific secondary antibody binding were also included. After washing 3 times with .1% TBST, sections were incubated with fluorophore-conjugated secondary antibody (raised against the primary antibody’s species) diluted in blocking buffer for 1 hour at room temperature in a humidified chamber. After washing, sections were counterstained with DAPI-containing mounting media (Vectashield, Cat no. H-1200-10) and imaged using a confocal (Nikon, A1RMP) microscope. Quantification was performed by manually counting the randomly selected fields and represented as positive cells per field.

### Isolation of lamina propria stromal cells from colon tissue

Mouse colon tissue was excised and opened longitudinally. The tissue was thoroughly washed three times in calcium and magnesium-free Hank’s Balanced Salt Solution (HBSS) to remove all fecal material. The colon was then cut into 2-5 cm pieces and incubated for 20 minutes on ice with a chelation buffer containing 30mM of EDTA in Ca++ and Mg++ free HBSS. Following incubation, the tissue was vigorously vortexed/shaken to detach epithelial cells, and the supernatant was discarded. Further, tissue pieces were washed with Ca++ and Mg++ free HBSS buffer until a clear supernatant, devoid of epithelial cells, was obtained. The Lamina Propria (LP) tissue pieces were then minced into the smallest possible fragments and incubated with 10 ml of pre-warmed dissociation buffer containing DMEM, 2-4 Wunsch units Liberase TM, 100µg/ml of DNase-I for 20 minutes at 37°C on a rocker. After incubation, tubes containing tissue were centrifuged at 300g for 5 minutes to pellet down the tissue pieces, while the supernatant was discarded. Further DMEM containing 10% FBS media was added to the tubes containing tissue and shaken vigorously to complete dissociation of the cells and then passed through a 70µm filter into a fresh tube. Finally, the cellular suspension was pelleted by centrifugation at 500g for 15 minutes at 4°C. For downstream immunophenotyping, the isolated cells were stained and sorted for myeloid cell populations using the following markers: PerCP-Cy5.5-CD45, BV786-CD11b, BV650-MHC II (M5/114.15.2), APC-F4/80, PE-Dazzle-CX3CR1, and PE-Cy7-Ly6C. Sorted cells were pelleted down and stored at – 80°C until further use.

### Quantitative Real-time PCR

qPCR was performed to determine the Cancer stem cells and β-Catenin target genes expression in colon tumor cells of Csf1r.iCre; Porcnfl/fl and Porcnfl/fl mice. Total RNA was isolated from tumor cells using the RNeasy micro/mini kit from Qiagen (Germantown, MD, USA) according to the manufacturer’s instructions. Briefly, cells were suspended in lysis buffer, and homogenization was performed for complete disruption of the cells. RNA was purified through a series of binding, washing, and elution steps using spin columns. The RNA concentration and purity were measured using a NanoDrop spectrometer (Thermo Scientific, Waltham, MA, USA). 1 µg of total RNA was reverse transcribed with the RNA to cDNA EcoDry™ Premix (Double Primed) (Takara Bio USA Inc., San Jose, CA, USA). The reaction was carried out for 1 h at 42°C, followed by a 10-minute. incubation at 70°C to end the reaction cycle. qPCR was carried out using the QuantStudio™ 7 Flex Real Time PCR System (Applied Biosystems™, New York, NY, USA) and SYBR Green Supermix (Bio-Rad, Hercules, CA, USA). β-Actin was used as the internal control for sample normalization. Relative gene expression was calculated using the ΔΔCt method. A complete list of primers used in this study is mentioned in Supplementary Table 1.

### HTB-38 cell culture

Human colorectal adenocarcinoma cell line (HTB-38) was purchased from ATCC. HTB-38 cells were cultured in McCoy’s 5A (modified) Medium containing 10% fetal calf serum and 1% penicillin/streptomycin. The cell cultures were maintained under standard cell culture conditions at 37°C and a humidity-controlled atmosphere with 5% CO2. Cells were passaged twice a week by splitting at a ratio of 1:3.

### Human Macrophage cultures

Human macrophages were cultured from PBMCs purchased from StemCell Technologies (Cat no. 70025.3). PBMCs were plated in T75 flasks. After 24 hrs of incubation, cells were transferred to a fresh tissue-culture flask and cultured for seven days with expansion media containing StemSpan™-XF with the addition of Expansion Supplement (StemCell Technology, Vancouver, BC, Canada) and 100 ng/ml of h-CSF (Cat No. 574808, BioLegend, San Diego, CA, USA). Following maturation, h-Mɸ were treated either with 20µM of Porcupine inhibitor-C59 (Cat no. 5148, Bio-Techne Corp., MN, USA) or 10µM of GSK-3β inhibitor-BIO (Cat no. S7198, Selleck Chemical, Houston, TX, USA) for 2 hours or in combination first with C59 treatment for 2 hours and then with BIO for 2 hour,s before use for the co-culture experiments.

### T cells culture

Human Pan T cells were purchased from StemCell Technologies (Cat no. 70024). Cells were cultured with T cells expansion media containing ImmunoCult™-XF T Cell Expansion Medium (Cat no. 10981; StemCell Technologies) with 10ng/ml of recombinant human-IL-2 (Cat no. 78036.3, StemCell Technologies). T cells were activated just before their use in the co-culture experiment using ImmunoCult™ Human CD3/CD28 T Cell Activator (Cat no. 10971, StemCell Technologies).

### Co-Culture assay

To study the interaction between tumor cells and immune cells (T cells and Human macrophages) on the expression of immune checkpoints, a co-culture assay was set up using a 6-well plate Transwell. The T cells (1.5 × 10^6 cells) were plated in the six-well plate per well, while the Cancer cells (1 × 10^6 cells) were embedded in the collagen Matrigel layer on the trans-well, and human macrophages (1.5 × 10^6 cells) were overlayed on top of the cancer cell-embedded collagen layer. The co-culture setup was incubated at standard cell culture conditions at 37°C with 5% CO2. After 48 hours of incubation, cells were harvested from the co-culture setup and analyzed for immune checkpoint expression using flow cytometry.

### Flow cytometry analysis

To investigate whether macrophage-derived WNT regulates immune checkpoint expression within the tumor microenvironment, colon tumor tissues were processed to isolate stromal immune cells for flow cytometric analysis. Following tissue dissociation and immune cell isolation, the resulting cell suspension was stained with specific antibodies to characterize key immune populations and their checkpoint profiles. Tissue-associated macrophages (TAMs) were identified using myeloid-specific markers: PerCP-Cy5.5-CD45, BV786-CD11b, BV650-MHC II (M5/114.15.2), APC-F4/80, PE-Dazzle-CX3CR1, and PE-Cy7-Ly6C. The proportion of myeloid-derived suppressor cells (MDSCs) within the total stromal immune compartment was also quantified. T cell populations, including pan T cells, cytotoxic T cells, and helper T cells, were identified using PerCP-Cy5.5-CD45, BV510-CD3, BV711-CD8, APC-CD25, PE-Cy7-CD4, and BV605-CD69. Colon cancer epithelial cells were identified based on EpCAM (CD326) expression. Further, Immune checkpoint expression (PD1 and PDL1) was analyzed across MDSCs, T cells, and tumor cells to determine potential regulatory interactions with macrophage-derived WNT signaling. Flow cytometry data were analyzed using FlowJo software (BD Biosciences, Franklin Lakes, NJ, USA).

### RNA sequencing

For RNA-seq analysis, sorted MDSCs from the colonic tumors were collected and immediately frozen at -80 °C to preserve RNA integrity. All the samples were shipped to BGI Global Genomics Services (BGI Hong Kong Tech Solution NGS Lab). BGI performed RNA extraction, quality assessment, mRNA purification, mRNA fragmentation, and cDNA reverse transcription. The TRIzol method was used to extract total RNA, and then a fragment analyzer was used for the assessment of quality, including measurements of the total RNA concentration, RNA integrity number (RIN) or RNA quality number (RQN), 28S/18S ratio, and fragment size. The integrity of the RNA was determined by agarose gel electrophoresis. Total RNA samples meeting the quality thresholds (RQN or RIN≥7.0, 28S/18S≥1.0) were selected (one sample in the NORMAL group was rejected).

The total RNA samples are first treated with DNase I to degrade any possible DNA contamination. Then the mRNA is enriched by using the oligo (dT) magnetic beads. Mixed with the fragmentation buffer, the mRNA is fragmented into short fragments. Then the first strand of cDNA is synthesized by using a random hexamer primer. Buffer, dNTPs, RNase H, and DNA polymerase I are added to synthesize the second strand. The double-strand cDNA is purified with magnetic beads. End reparation and 3’-end single nucleotide A (adenine) addition is then performed. Finally, sequencing adaptors are ligated to the fragments. The fragments are enriched by PCR amplification. During the QC step, Agilent 2100 Bioanalyzer and ABI StepOnePlus Real-Time PCR System are used to qualify and quantify the sample library. The library products are ready for sequencing via Illumina HiSeqTM 2000 or another sequencer when necessary. Libraries were then sequenced on an Illumina NovaSeq 6000 sequencing machine (Illumina, San Diego, CA) with a strand-specific, 100-cycle paired-end resolution. After the library was constructed and its quality was verified, the single-stranded circular DNA molecules were copied by rolling circle amplification to form DNA nanoballs (DNBs). A sequence read length of 150 bp was obtained by sequencing on the BGISEQ-500 platform powered by combinatorial probe-anchor synthesis (cPAS). The Data was finally visualized by Mr. Tom’s analysis platform.

### Transcriptomic Digital Spatial Profiling

The GeoMx Digital Spatial Profiling (DSP) platform (Bruker NanoString Technologies) was used to characterize the tumor biology of mouse colon villi and crypts with a focus on uncovering the transcriptomic factors that contribute to tumor formation following radiation/treatment. Spatial transcriptomic profiling was performed using the mouse Whole Transcriptome Atlas (WTA) assay, which measures the expression of approximately 21,000 genes associated with tumor biology. The assay uses a cocktail of in situ hybridization (ISH) probes that bind to the mRNA targets within the fixed tissue sections. The ISH probes are conjugated to light-sensitive, photocleavable DNA-barcoded oligos, which provide highly multiplexed transcriptomic data. Fluorescently labeled antibodies are used to elucidate the cell types of interest within the tissue sections for analysis. DSP technology and the GeoMx platform have been extensively used and reviewed since their inception for studies in a variety of cancer types (PMID: 38850363; PMID: 35847846; PMID: 38378868; PMID: 37106610

A total of 5 colon tissue samples (n=5/group) from *Porcn^fl/fl^* (wild type) and *Csf1r.iCre; Porcn^fl/fl^* (WNT Null) mice were processed and analyzed in this study. For each sample, ∼ 5 regions of interest (ROIs) were selected for DSP analysis. ROIs were further segmented as described below. DSP of the colon tissue samples from mice was performed according to NanoString’s GeoMx protocols for manual slide preparation, hybridization, staining, collection, quantification, and data normalization. Briefly, this entails using an unstained slide for each patient sample containing a 5 µM FFPE section of the tissue for overnight hybridization using a cocktail of the in-situ hybridization (ISH) probes. Following overnight hybridization, the slides were stained with fluorescently conjugated antibodies against PanCK (epithelial cell marker), CD68 (macrophage marker), CD3 (T-cell marker), and a SYTO-13 dye (nuclear stain to visualize nuclei). The hybridized and fluorescently stained slides were then scanned on the GeoMx instrument, and ROIs for analysis were selected. ROIs were segmented based on the staining pattern of the morphological markers (PanCK, CD68, and CD3). The photocleavable DNA barcoded-oligos from each segment were collected into individual wells of a 96-well plate. Once all tissue samples were processed and collected, next-generation sequencing (NGS)-based quantification was performed. NGS libraries were sequenced using the Illumina NextSeq 550 or NextSeq 2000 instruments. Illumina FASTQ sequencing files were processed using the GeoMx NGS Pipeline version 2.3.3.10 to generate digital count conversion (DCC) files, which were subsequently analyzed using the GeoMx DSP Analysis Suite version 3.0 to perform data QC and Q3 normalization using all target groups.

### Statistical and Bioinformatics analysis

Mice were randomly assigned to each experimental and control group after genotyping. A minimum of 10 mice per group (n=10) were used for the study, and all animal studies were performed three times. For colon sampling, regions were randomly selected for digital image acquisition and subsequent quantitation. Digital image data was evaluated in a blinded manner with respect to treatment. A two-sided Student’s t-test was employed to determine significant differences between experimental groups. All data are presented as the mean ± standard deviation (SD). Differences between groups with p < 0.05 were considered statistically significant. RNA Seq Primary sequencing data produced by Illumina HiSeqTM 2000, called raw reads, are subjected to quality control (QC) to determine if a resequencing step is needed. After QC, raw reads are filtered into clean reads, which are aligned to the reference sequences. QC of alignment was performed to determine if resequencing is needed. The alignment data were utilized to calculate the distribution of reads on reference genes and the mapping ratio. Once alignment result passes QC, the downstream analysis is done, which includes gene expression and deep analysis based on gene expression (PCA/correlation/screening differentially expressed genes, and so on). Further, deep analysis was done based on DEGs, including Gene Ontology (GO)enrichment analysis, KEGG pathway enrichment analysis, cluster analysis, protein-protein interaction network analysis, and finding transcription factors. The normalized gene expression data in the GeoMx study analyses were used using the Bioconductor Package in R. The package provides functions for reading in DCC and PKC files based on an Expression Set-derived object. Normalization and QC functions are also included.

## Results

### Inhibition of macrophage derived WNT release promote colonic tumor

To investigate the role of macrophage-derived Wnt in a mouse model of colon cancer, *Csf1r.iCre;Porcn^fl/fl^* mice, which exhibit a macrophage-specific deletion of Porcn [9], were utilized (Figure 1A). Deletion of porcupine inhibits the release of physiological Wnt from macrophages. Following AOM/DSS treatment, *Csf1r.iCre;Porcn^fl/fl^* mice developed a significantly higher number of colonic macroscopic and microscopic tumors compared to wild-type littermates (*Porcn^fl/fl^*). Multiple parameters associated with AOM/DSS-induced colon cancer, including Disease Activity Index (DAI) score (Figure 1 B), tumor counts (Figure 1 C-F), representation of tumor in colon by colonoscopy and transmission electron microscopy (Figure 1 G, H), and the presence of cytokeratin (CK) antigen (Figure 1 I, J), indicated that *Csf1r.iCre;Porcn^fl/fl^* mice were more susceptible to AOM/DSS colon cancer than Porcn*^fl/fl^* mice. Further analysis of plasma cytokine from *Csf1r.iCre;Porcn^fl/fl^* and *Porcn^fl/fl^* mice demonstrated significantly low anti-inflammatory (Figure 1K) and higher inflammatory (Figure 1L) cytokine expression in *Csf1r.iCre;Porcn^fl/fl^* compared to *Porcn^fl/fl^* mice. These findings suggest that the absence of macrophage-derived Wnt promotes colitis associated colon cancer in mice.

**Figure 1:**
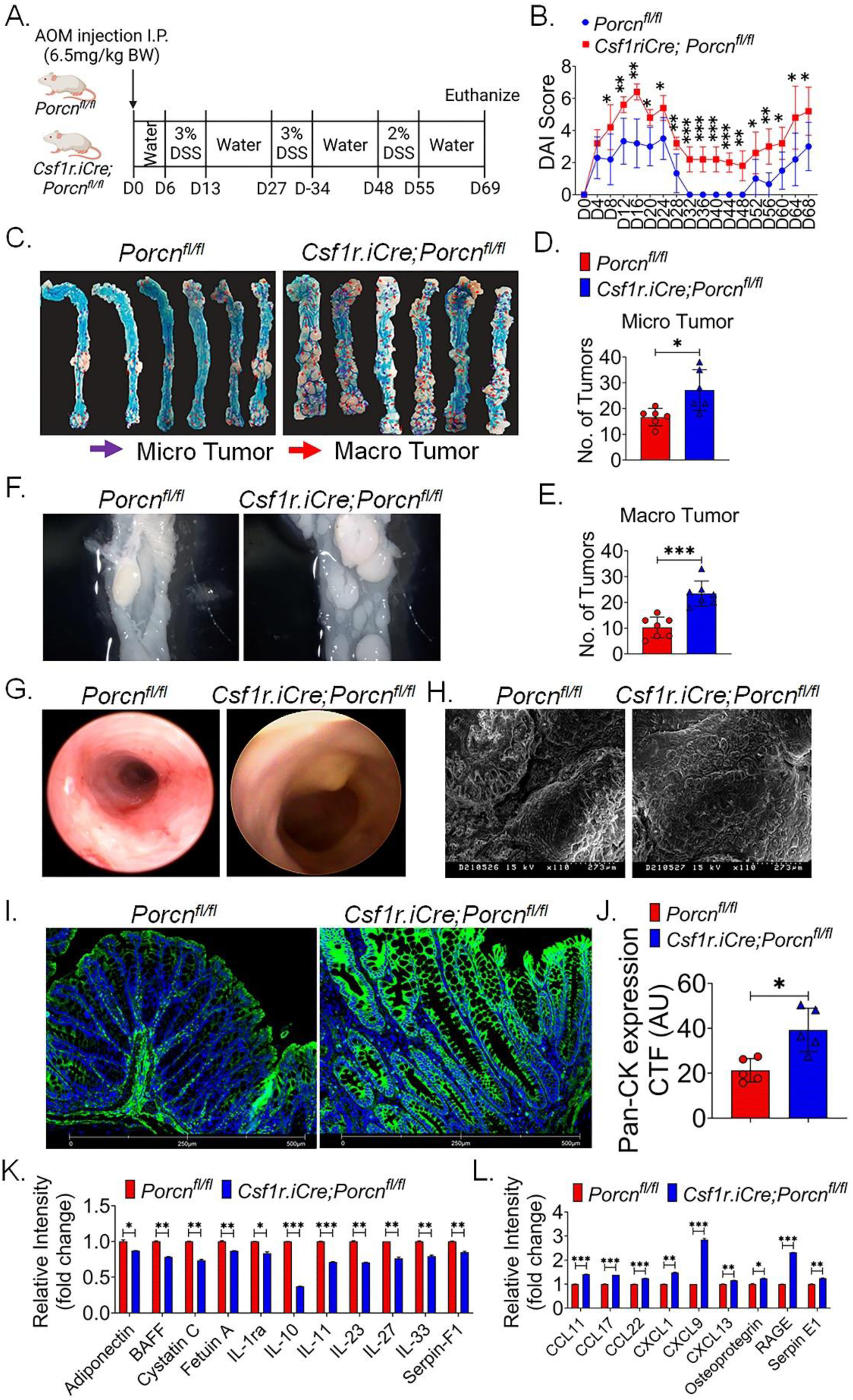
Csf1r.iCre;Porcn^fl/fl^ mice are sensitive to AOMDSS induced colon cancer. A. Schematic diagram of AOMDSS treatment in Csf1r.iCre;Porcn^fl/fl^ and wild type (WT) mice. B. DAI scored over 68 days since initiation of AOMDSS treatment. Csf1r.iCre;Porcn^fl/fl^ demonstrated significantly high DAI score compared to WT mice. C. Photographic images demonstrate higher presence of macro and micro tumor in Csf1r.iCre;Porcn^fl/fl^ mice compared to WT mice at 69 days post AOMDSS treatment initiation. D-E. Histogram demonstrates significantly higher presence of macro and micro tumors in Csf1r.iCre;Porcn^fl/fl^ mice compared to WT mice. F. Dissection macroscopic images (10X) of AOMDSS colon tumor demonstrating higher number in Csf1r.iCre;Porcn^fl/fl^ mice compared to WT mice G. Colonoscopy demonstrating presence of tumor in colonic luminal wall in Csf1r.iCre;Porcn^fl/fl^ mice at 55 days post AOMDSS treatment initiation. H. Scanning Electron Microscopic image in WT and Csf1r.iCre;Porcn^fl/fl^ mice. Microscopic images showing higher presence of Pan CK epithelial cell marker in Csf1r.iCre;Porcn^fl/fl^ mice colon tumor compared to WT mice. J. Histogram demonstrated significantly higher Pan CK expression in Csf1r.iCre;Porcn^fl/fl^ mice colon tumor tissue compared to WT mice. K-L. Plasma cytokine analysis demonstrated significantly downregulation of anti-inflammatory cytokine and higher presence of inflammatory cytokine and in Csf1r.iCre;Porcn^fl/fl^ mice compared to WT mice. Data presented as the mean ± SD. (Significant level, *: p<0.05; **: p<0.005; ***: p<0.0005).

### Inhibition of macrophage derived WNT release induces stem cell population in colon cancer

Consistent with the observed a higher tumor burden in *Csf1r.iCre;Porcn^fl/fl^* mice as observed by H&E staining (Figure 2 A), and an increased number of Ki67-positive cells were found (Figure 2 B), suggesting heightened proliferation of colonic malignant cells. Both Lgr5+ [10] and DCLK1 [11] cells are considered stem/progenitor populations in colon cancer. The AOM/DSS model demonstrated that *Csf1r.iCre;Porcn^fl/fl^* mice showed a greater enrichment of Lgr5+ and DCLK1 cells (Figure 2 C) compared to wild-type (WT) mice, supporting the observation of higher tumor burden in mice with *Porcn* null macrophages. Furthermore, an increase in WNT target gene expression was observed (Figure 2 D) within the colon tumor cells of *Csf1r.iCre;Porcn^fl/fl^* mice, correlating with a higher number of colon cancer stem cells (Figure 2 E).

**Figure 2.**
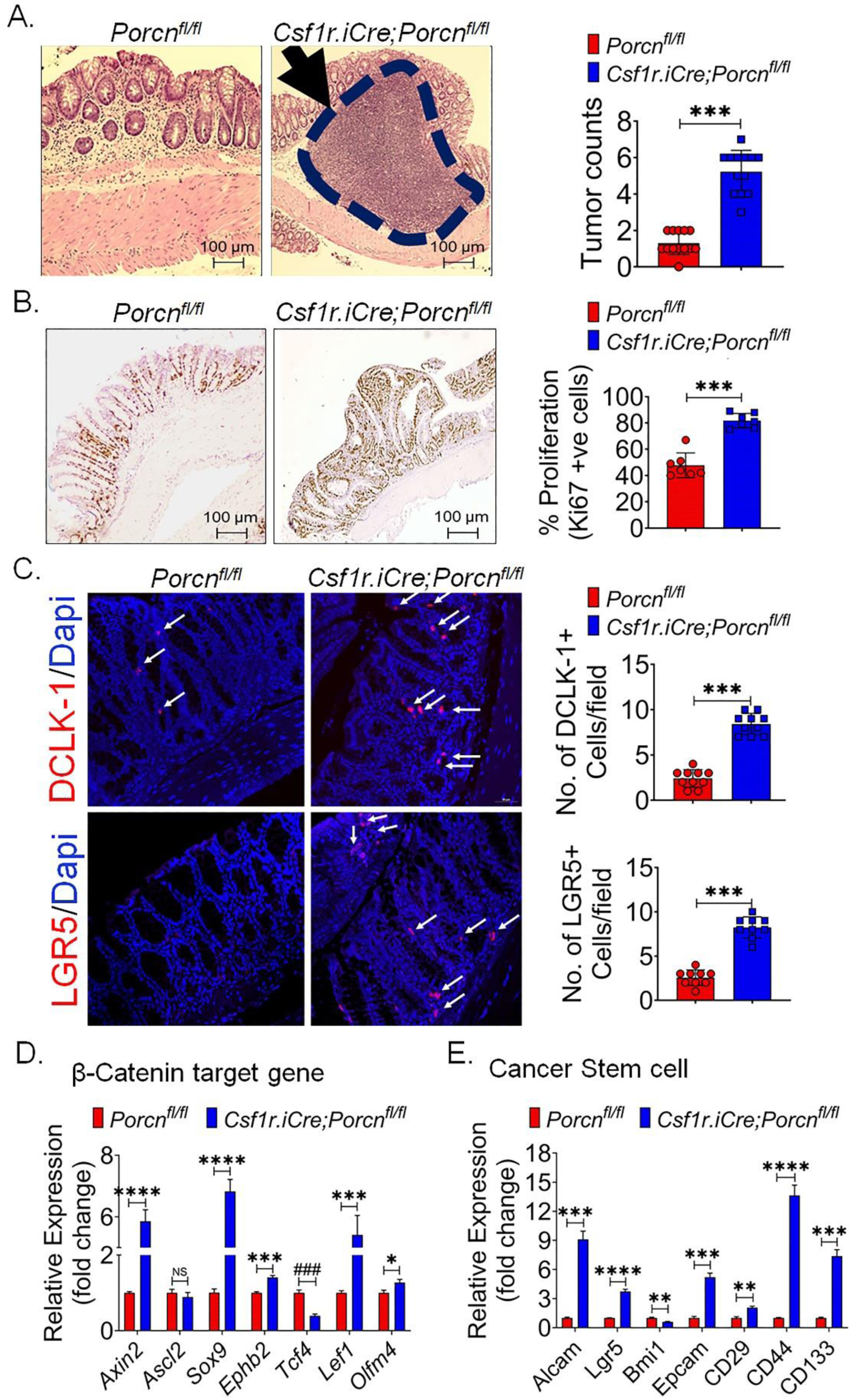
Csf1r.iCre;Porcn^fl/fl^ mice demonstrated higher proliferation of colon cancer cells with enrichment of cancer stem cells following AOMDSS treatment. A. Microscopic imaging along with Histogram demonstrated presence of higher number of tumors in colonic epithelium in Csf1r.iCre;Porcn^fl/fl^ mice compared to WT mice. B. Microscopic imaging along with Histogram demonstrated presence of higher number of Ki67+ve cells in colonic epithelium in Csf1r.iCre;Porcn^fl/fl^ mice compared to WT mice. C. Confocal microscopic imaging along with histogram demonstrating presence of higher number of DCLK-1 positive (upper panel, stained red & indicated with arrow) and Lgr5 positive (lower panel, indicated stained red & indicated with arrow) cancer stem cells in Csf1r.iCre;Porcn^fl/fl^ mice compared to WT mice. D. qPCR analysis of mRNA expression for β-Catenin target genes demonstrated higher expression in Csf1r.iCre;Porcn^fl/fl^ mice compared to WT mice. E. qPCR analysis of mRNA expression for cancer stem cell marker demonstrated higher expression in Csf1r.iCre;Porcn^fl/fl^ mice compared to WT mice. Data presented as the mean ± SD. (Significant level, *: p<0.05; **: p<0.005; ***: p<0.0005; NS: not significant).

### Absence of macrophage derived WNT alters checkpoint expression in macrophages

Tumor-associated macrophages (TAMs) are key players in fostering an immunosuppressive tumor microenvironment [4, 12, 13]. Unlike macrophages in acute inflammation, which phagocytose and present antigens to stimulate adaptive immunity, TAMs inhibit effector T and NK cell activity through soluble immunosuppressive factors and membrane-bound immune checkpoint molecules, such as Programmed Death-Ligand 1 (PD-L1) [14–16]. Paracrine signals from cancer cells reprogram TAMs to adopt an immunosuppressive and tumor-promoting phenotype [14].

Our study revealed that the absence of macrophage-derived WNT in *Csf1r.iCre;Porcn^fl/fl^* mice enhances PD-1 and PD-L1 expression in TAMs. Flow cytometry of AOMDSS colon tumor cells showed a higher number of PD-1 and PD-L1 expressing Ly6C-Cx3CR1+ TAMs in *Csf1r.iCre;Porcn^fl/fl^* mice compared to wild-type (WT) mice (Figure 3 A-C). Immunofluorescence staining of AOMDSS colon tumor tissue further confirmed an increased presence of PD-1 (Figure 3 D, E) and PD-L1 (Figure 3 F, G) expressing F4/80+ macrophages. The regulation of immune checkpoint expression, including PD-1 and PD-L1, in macrophages remains an area that has not been thoroughly investigated.

**Figure 3.**
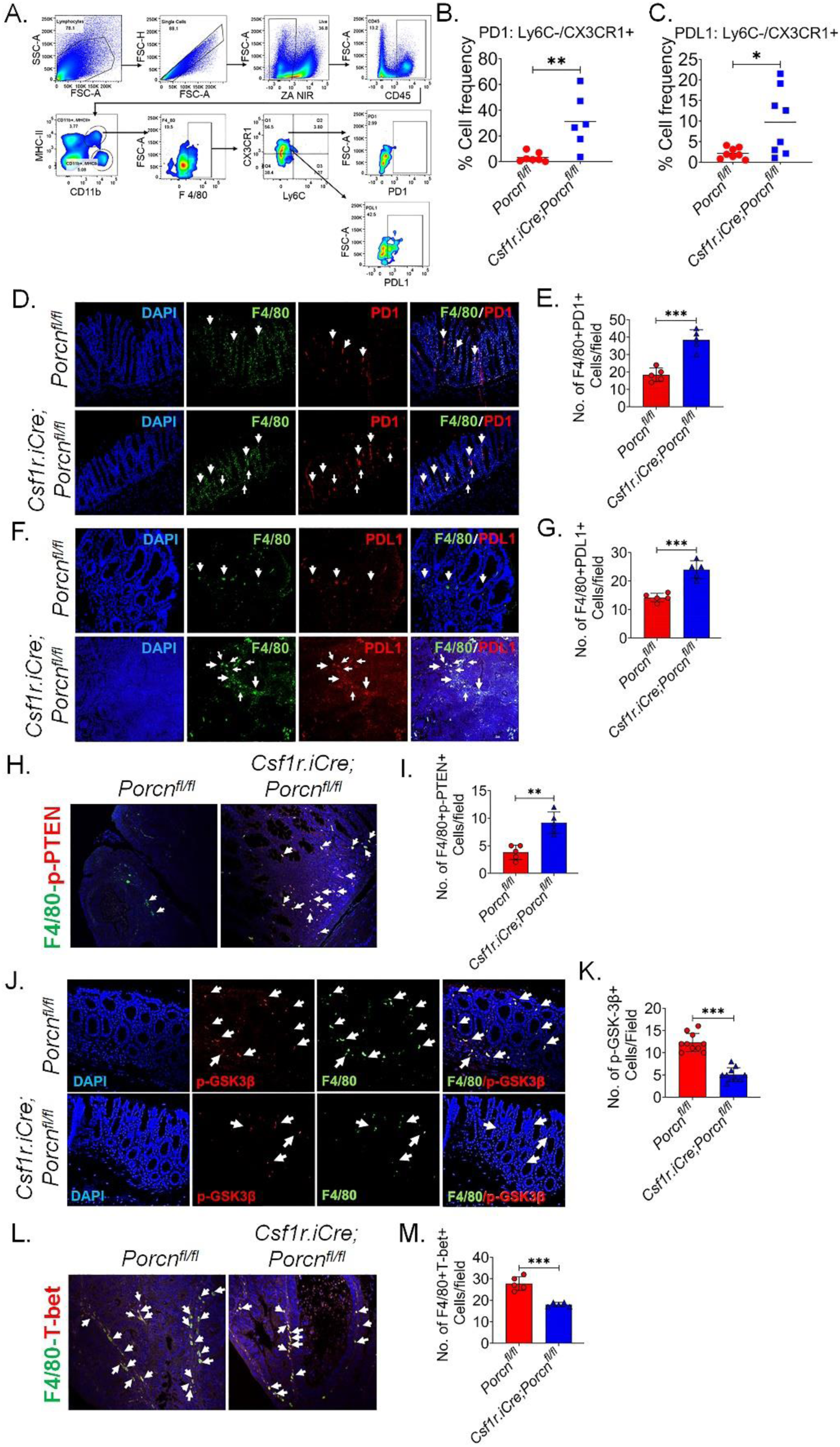
Tumor associated macrophages (TAM) are more immunosuppressive in Csf1r.iCre;Porcn^fl/fl^ mice colon tumor compared to WT mice. A. Flowcytometry plot demonstrates gating strategy to achieve PD1 and PDL1 positive TAM in colon tumor cells. B, C. Percent of PD1 and PDL1 positive TAMs (Ly6C-Cx3CR1+) are much higher in Csf1r.iCre;Porcn^fl/fl^ mice colon tumor compared to WT mice. D. Confocal microscopic images demonstrate higher presence of PD1+ve macrophages (F480+ve) in Csf1r.iCre;Porcn^fl/fl^ mice colon tumor tissue compared to WT mice. F480+ve cells were green, PD1+ cells were Red and F480+PD1+ cells were identified by pink color (white arrow). E. Histogram demonstrating quantification of F480+PD1+ cells. Csf1r.iCre;Porcn^fl/fl^ mice colon tumor tissue sections showed significantly higher number of F480+PD1+ cells compared to WT mice. F. Confocal microscopic images demonstrating higher presence of PDL1+ve macrophages (F480+ve) in Csf1r.iCre;Porcn^fl/fl^ mice colon tumor tissue compared to WT mice. F480+ve cells were green, PDL1+ cells were Red and F480+PDL1+ cells were identified by pink color (white arrow). G. Histogram demonstrating quantification of F480+PDL1+ cells. Csf1r.iCre;Porcn^fl/fl^ mice colon tumor tissue sections showed significantly higher number of F480+PDL1+ cells compared to WT mice. H. Confocal microscopic images demonstrating presence of p-PTEN +ve macrophages (F480+ve) in Csf1r.iCre;Porcn^fl/fl^ mice colon tumor tissue compared to WT mice. F480+p-PTEN +ve cells were identified by white arrow. I. Histogram demonstrating quantification of F480+ p-PTEN +ve cells. Csf1r.iCre;Porcn^fl/fl^ mice colon tumor tissue sections showed significantly higher number of F480+ p-PTEN +ve cells compared to WT mice. J. Confocal microscopic images demonstrate less presence of F480+ pGSK3beta+ve macrophages in Csf1r.iCre;Porcn^fl/fl^ mice colon tumor tissue compared to WT mice. F480+ pGSK3beta+ve cells were identified with white arrow. K. Histogram demonstrating quantification of F480+ pGSK3beta+ve cells. Csf1r.iCre;Porcn^fl/fl^ mice colon tumor tissue sections showed significantly a smaller number of F480+ pGSK3beta+ve macrophages compared to WT mice. L. Confocal microscopic images demonstrate less presence of T-bet +ve macrophages (F480+ve) in Csf1r.iCre;Porcn^fl/fl^ mice colon tumor tissue compared to WT mice. F480+T-bet +ve cells were identified by white arrow. M. Histogram demonstrating quantification of F480+T-bet+ cells. Csf1r.iCre;Porcn^fl/fl^ mice colon tumor tissue sections showed significantly lower number of F480+ T-bet+ cells compared to WT mice. Data presented as the mean ± SD. (Significant level, *: p<0.05; **: p<0.005; ***: p<0.0005).

Previous research has indicated that PTEN, a tumor suppressor gene, dephosphorylates the lipid signaling intermediate PIP3, thereby inhibiting AKT activity, a key effector of the PI3K signaling pathway [17, 18]. Loss of PTEN leads to the activation of the PI3K/AKT pathway, resulting in increased PD-L1 expression. PTEN activity is also regulated post-translationally through phosphorylation, which can inhibit its function. Immunofluorescent staining of colon tumor tissue in *Csf1r.iCre;Porcn^fl/fl^* mice demonstrated a significant upregulation of phospho-PTEN positive macrophages compared to WT mice (Figure 3 H, I).

GSK3β, a multifunctional serine/threonine kinase, plays a crucial role in regulating PTEN activity. Phosphorylation of PTEN by GSK3β at specific sites, such as Ser-362 and Thr-366, inhibits its activity. Conversely, inhibition of GSK3β activity, often through phosphorylation, decreases PTEN phosphorylation and restores PTEN activity [19, 20]. Immunofluorescence staining of colon tumor tissue macrophages from *Csf1r.iCre;Porcnfl/fl* mice showed a significant reduction in pGSK3β levels compared to wild-type mice (Figure 3 J, K). These observations suggest that macrophages in *Csf1r.iCre;Porcnfl/fl* mice fail to inhibit GSK3β activity, leading to increased GSK3β-mediated PTEN phosphorylation and inhibition. WNT ligand-receptor interactions are known to phosphorylate and inhibit GSK3β. Given that macrophages express WNT receptors [21], our study suggests the presence of a WNT autofeedback loop within colon tumor macrophages that typically restores PTEN activity. However, the absence of WNT in *Csf1r.iCre;Porcnfl/fl* macrophages disrupts this autofeedback loop, resulting in sustained GSK3β activity, PTEN inhibition, and the induction of PDL1 expression in these tumor-associated macrophages (TAMs).

Inhibition of GSK3β has been shown to reduce PD-1 transcription by upregulating TBET expression [22]. Given the observation of increased PD-1 expression in tumor-associated macrophages (TAMs) from *Csf1r.iCre;Porcn^fl/f^l* mice, the levels of TBET in these TAMs were investigated. Immunofluorescence staining of colon tumors revealed significantly lower TBET expression in TAMs from *Csf1r.iCre;Porcn^fl/fl^* mice compared to wild-type controls (Figure 3 L, M). This finding suggests that the absence of WNT signaling in *Csf1r.iCre;Porcn^fl/fl^* TAMs disrupts a crucial autofeedback loop, leading to the reversal of GSK3β activation, reduced TBET expression, and subsequent upregulation of PD-1.

### Pharmacological inhibition of GSK3β in macrophages reduces colon tumor burden in AOMDSS treated mice

To validate the role of GSK3β in reducing colon tumors, pharmacological targeting of GSK3β was performed (Figure 4 A). Delivery of a macrophage-targeted liposomal GSK3β inhibitor during AOMDSS treatment significantly reduced the tumor burden in *Csf1r.iCre;Porcn^fl/fl^* mice (Figure 4 B-D). These results strongly support the conclusion regarding the WNT autofeedback loop in macrophages. Pharmacological targeting of GSK3β reversed the dependence of tumor-associated macrophages (TAMs) on the WNT autofeedback loop to overcome their immunosuppressive phenotype. Additionally, GSK3β inhibitor treatment also reduced tumor burden in WT mice (Figure 4 B-D), which suggests that additional inhibition of GSK3β, even in the presence of a WNT autofeedback loop, promotes tumor reduction.

**Figure 4.**
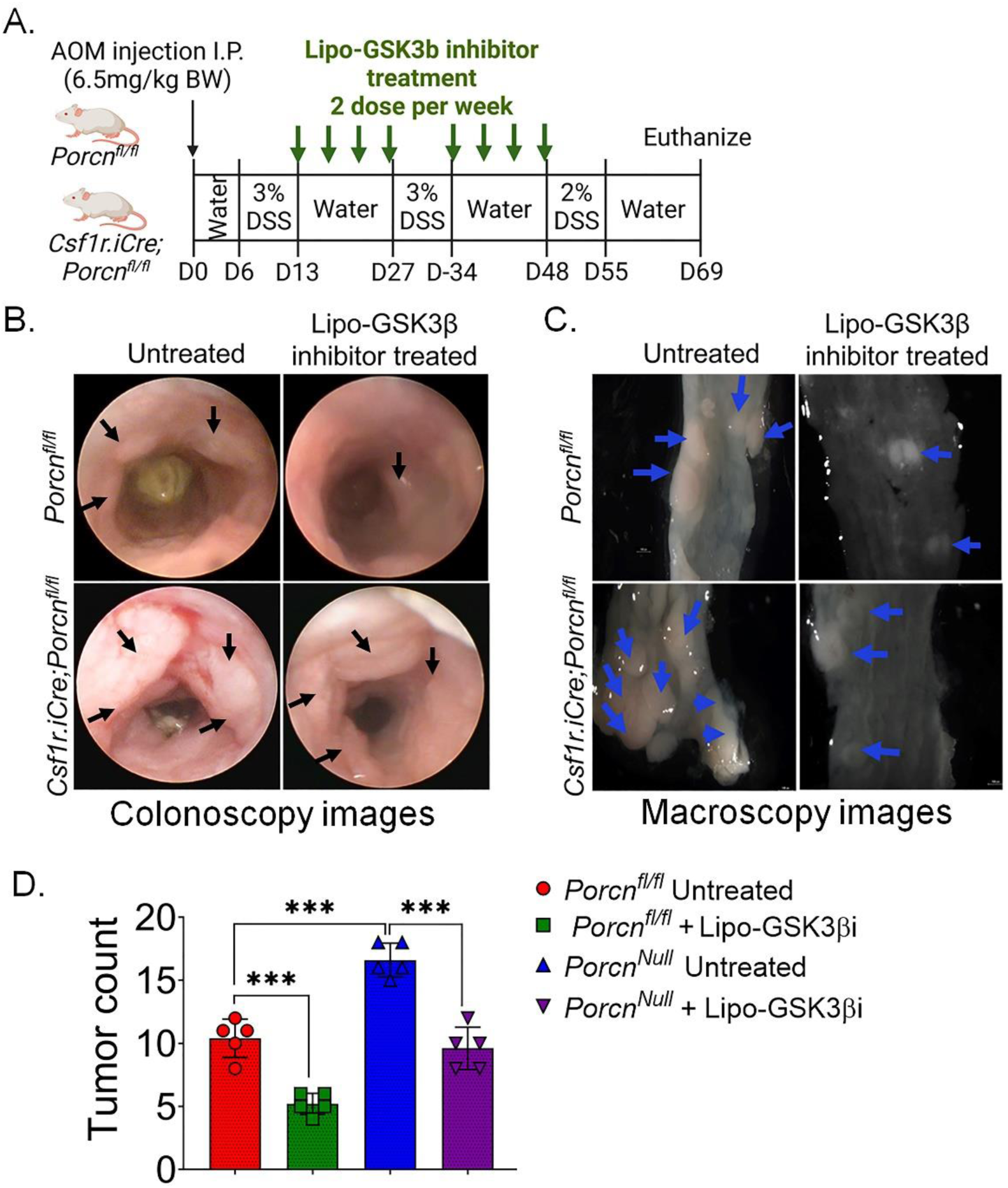
Macrophage specific inhibition of GSK3β modulates AOMDSS induced tumor growth. A. Schematic diagram for AOMDSS and Lipo-Gsk3β inhibitor treatment in mice. B. Colonoscopy images of AOMDSS colon tumor (indicated with arrow). Treatment with Lipo-Gsk3β reduces the AOMDSS tumor burden in Csf1r.iCre;Porcn^fl/fl^ mice. C. Dissection macroscopic images (10X) of AOMDSS colon tumor (indicated with arrow). Treatment with Lipo-Gsk3β reduces the AOMDSS tumor burden in Csf1r.iCre;Porcn^fl/fl^ mice. D. Histogram demonstrating significant downregulation of AOMDSS induced tumor burden with Lipo-Gsk3β inhibitor in both Csf1r.iCre;Porcn^fl/fl^ and WT mice. Data presented as the mean ± SD. (Significant level, ***: p<0.0005).

### Absence of macrophage derived WNT promotes immunosuppressive microenvironment in AOMDSS colon tumor

Transcriptomic analysis of colon tumor tissue using GeoMx revealed that loss of macrophage-derived WNT reprograms the immune microenvironment of AOMDSS induced colon tumors. In *Csf1r.iCre; Porcn^fl/fl^* mice, CD3⁺ T cells exhibited transcriptional alterations that favored immune evasion, including upregulation of *C5ar1*, *Rara*, and *Casp4*, while downregulation of the protective gene *Smcr8* (Figure 5 A). Activation of *C5ar1* on TAMs and MDSCs promotes recruitment of suppressive myeloid cells and secretion of IL-10 and TGF-β, directly dampening cytotoxic T cell activity and facilitating tumor progression [23]. Aberrant *Rara* signaling drives CD4⁺ T cells toward dysfunctional effector subsets that fail to support antitumor immunity, promoting a pro-tumor immunosuppressive environment [24], while elevated Casp4 activates inflammasomes that recruit suppressive myeloid cells and further weaken cytotoxic T cell responses, facilitating tumor immune evasion [25, 26]. In contrast, *Smcr8* normally contains TLR signaling to maintain immune homeostasis and effective antitumor surveillance, and its downregulation further enhances immune suppression [27]. CD68⁺ macrophages were also transcriptionally reprogrammed, exhibiting enrichment of *Trim31* and *Ccl12* along with elevated *Ccr1* and *Tlr4* expression, consistent with a pro-tumor immunosuppressive phenotype. *Trim31* integrates NF-κB, p53, and WNT/β-catenin signaling to reinforce macrophage-mediated immune suppression [28], *Ccl12* recruits monocytes that differentiate into TAMs inhibiting cytotoxic T cell function and promoting angiogenesis [29], *Ccr1* drives M2-like polarization, and chronic *Tlr4* activation sustains production of immunosuppressive cytokines and pro-angiogenic factors, collectively supporting tumor growth [30, 31]. In parallel, EpCAM⁺ tumor epithelial cells upregulated *Irf11*, whose dysregulation impairs TLR- and type I interferon-driven antitumor pathways, reduces epithelial immune surveillance, and favors tumor immune escape [32]. Collectively, these data demonstrate that deletion of *Porcn* in macrophages demolishes WNT-dependent immune regulation, driving multicellular transcriptional reprogramming across T cells, macrophages, and epithelial cells that establishes a pro-tumor, immunosuppressive microenvironment in colon tumors.

**Figure 5.**
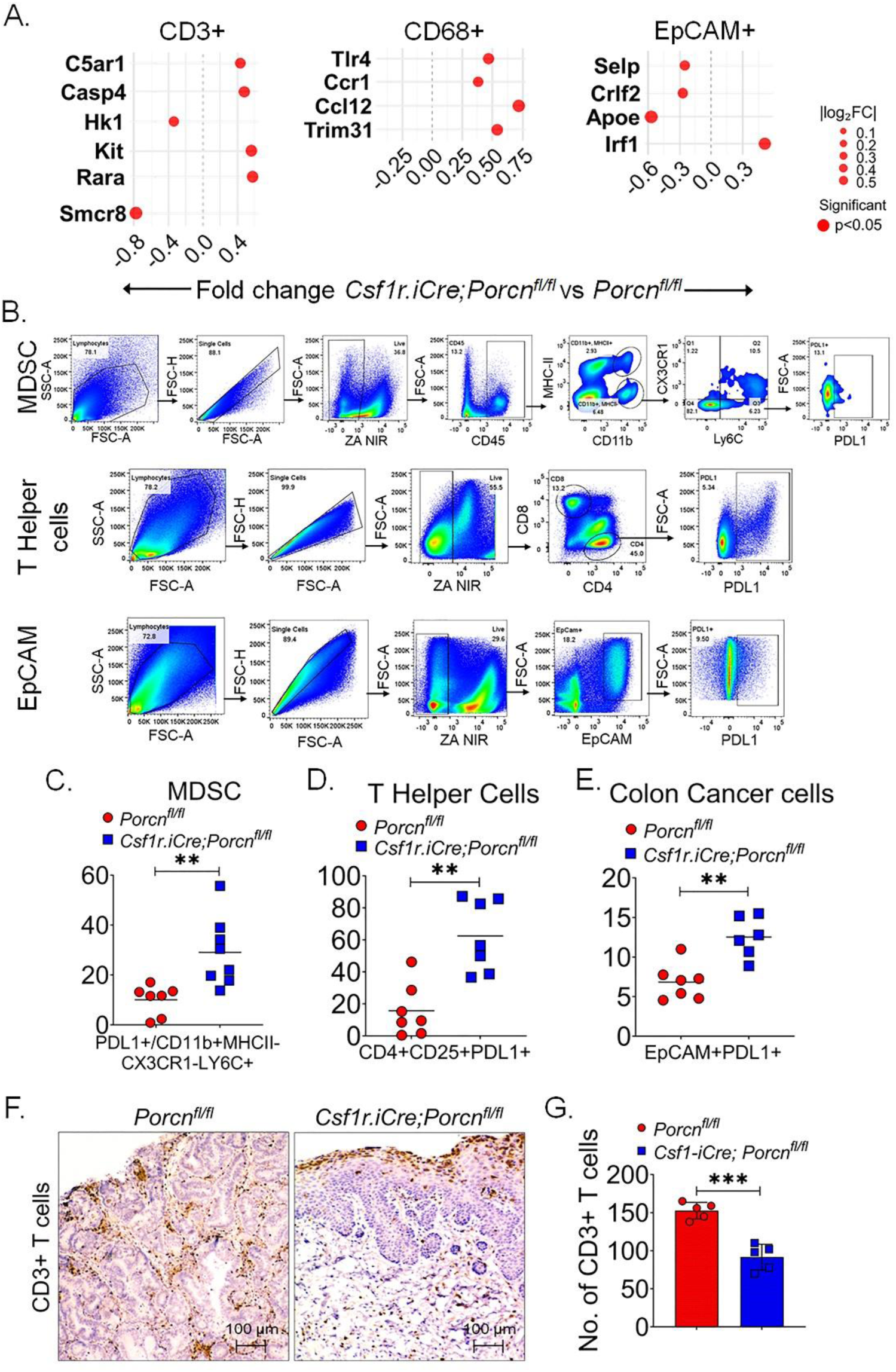
Csf1r.iCre;Porcn^fl/fl^ mice demonstrated immunosuppressive tumor microenvironment in AOMDSS induced colon tumor compared WT mice. A. GeoMx transcriptomic analysis on FFPE colon tumor sections demonstrated higher expression of immunosuppressive genes in CD3+ T cells, CD68+ macrophages and Epcam+ epithelial cells in Csf1r.iCre;Porcn^fl/fl^ mice compared to WT mice. B. Flowcytometry plot demonstrating gating strategy to achieve PDL1 positive MDSCs, T helper cells and Cancer cells. C-E. Dot plot demonstrates higher PDL1 expressing MDSCs (C), T helper cells (D), and Cancer cells (E). F, G. Immunohistochemistry demonstrates lower CD3+ T cell infiltration in Csf1r.iCre;Porcn^fl/fl^ mice colon tumor compared to WT mice. At least ten randomly selected high-power fields per section were counted and expressed as number of positive cells per total counted cells per field. Data presented as the mean ± SD. (Significant level, **: p<0.005).

To examine the involvement of macrophage derived WNT on the expression of PDL1 in other immune cells, including T cells and immunosuppressive MDSCs, flow cytometry was used on cells from the tumor microenvironment. Research has shown that PI3K signaling in macrophages affects PDL1 expression in both macrophages and T cells. Additionally, macrophages are known to regulate immunosuppressive signals in MDSCs [33]. The results in the AOMDSS model showed that PDL1 expression in T cells, MDSCs, and colon cancer cells were upregulated in *Csf1r.iCre;Porcn^fl/fl^* mice compared to WT mice (Figure 5 B-E). RNA seq analysis followed by KEGG pathway enrichment analysis of sorted MDSCs from the colonic tumors demonstrates significant upregulation of PI3K-AKT pathway involved in upregulation of PDL1 expression [34] (Supplementary Figure 1). Immunohistochemistry of colon tumor tissue demonstrated significantly lower number of CD3+ T cells in *Csf1r.iCre;Porcn^fl/fl^* mice compared to WT mice (Figure 5 F, G). This result further supports the role of WNT signaling in macrophages to overcome immunosuppressive tumor microenvironment.

To determine the translational relevance of these observations, an *ex-vivo* co-culture study was performed using human macrophages treated with or without the Porcupine inhibitor C59, human colorectal cancer cells (COLO205), and human T cells (Figure 6 A). Flow cytometry of cells from these co-cultures showed that C59-mediated inhibition of Porcupine in human macrophages increased PDL1 expression in all three cell types (CD4, CD8 and macrophages) compared to co-cultures with untreated macrophages (Figure 6 B-E). However, treating C59-exposed macrophages with a GSK3beta inhibitor before co-culture with COLO205 and human T cells reversed this PDL1 increase (Figure 6 B-E). These findings further suggest that the inhibition of GSK3β in human macrophages is important to overcome immunosuppressive microenvironment in human colon cancer.

**Figure 6.**
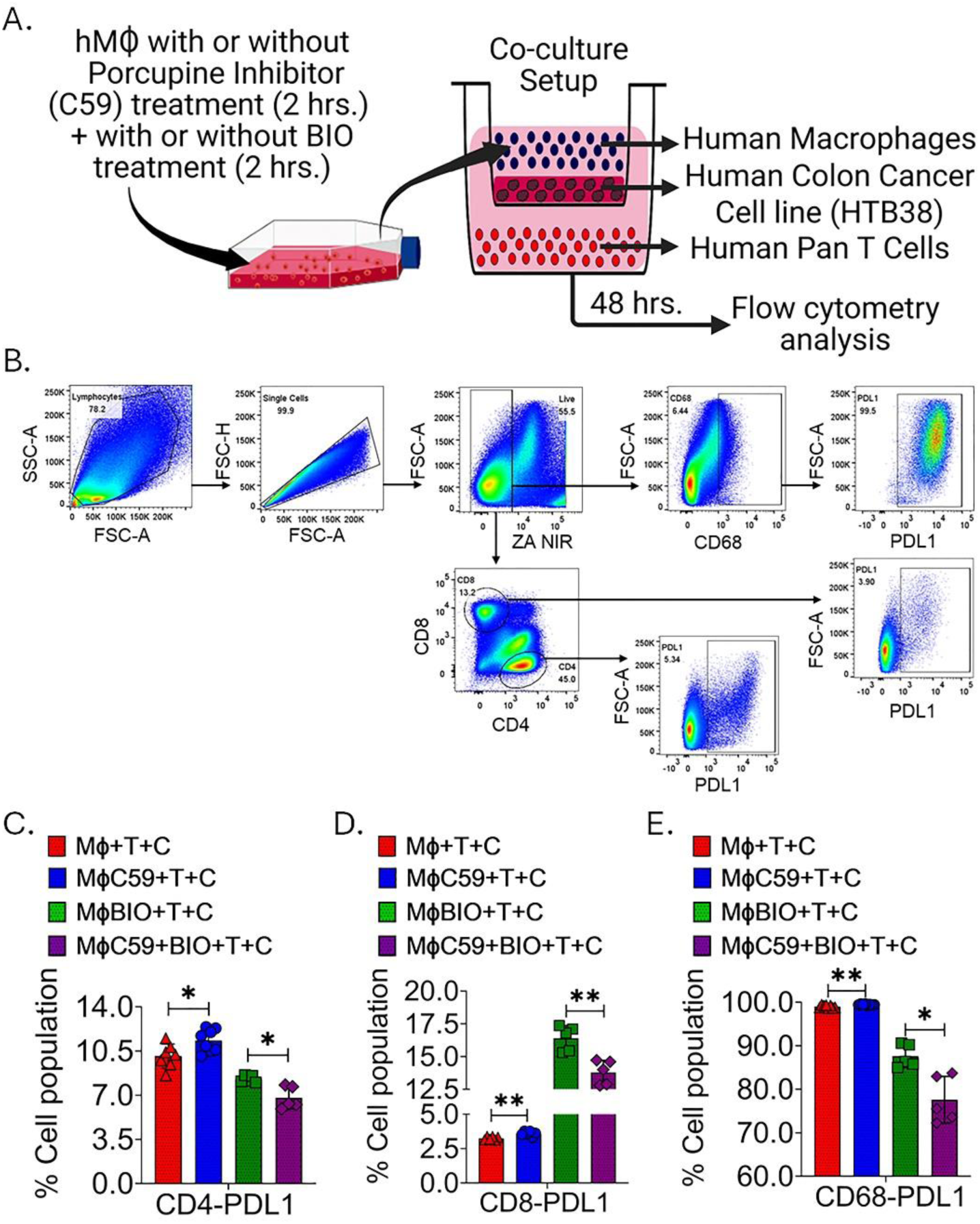
Pharmacological Inhibition of Porcupine in human macrophages alters PDL1 expression in T cells and CD68+ macrophages in a co-culture of human colon cancer cells COLO205, T cells and Macrophages. A. Schematic diagram of ex-vivo co-culture system consisting Human colon cancer cells COLO205, Human macrophages with/without Porcupine inhibitor C59 and/or BIO (GSK3β inhibitor) and human pan T cells. B. Flowcytometry plot demonstrating gating strategy to determine CD68 cells, CD4 cells and CD8 cells with PDL1 expression. C-E. Histogram demonstrating PDL1 expression in CD4 cells (C), CD8 cells (D), and CD68 cells (E). C59 treatment in human macrophages increases PDL1 expression in CD4 cells, CD8 cells and CD68 cells. However, treatment with BIO reduces PDL1 expression in CD4 cells, CD8 cells and CD68 cells. Data presented as the mean ± SD. (Significant level, *: p<0.05; **: p<0.005).

## Discussion

Colorectal cancer (CRC) is characterized by an immunosuppressive tumor microenvironment (TME) that significantly limits effective therapeutic interventions [35]. Macrophages within the TME play a crucial role in promoting and regulating this immunosuppression [36]. While macrophages have been a therapeutic target for controlling tumor initiation and growth for many years, the recent identification of distinct macrophage phenotypes with critical functions in the TME has led to preclinical studies addressing these phenotypes to overcome tumor burden [37, 38]. However, paracrine signals released from these macrophages also play a major role in tumor initiation and progression [39, 40], and therapeutic targeting of specific macrophage phenotypes has achieved limited success [41, 42] due to overlapping functions that either promote or inhibit tumor progression. Therefore, identifying the key macrophage-derived signals influencing tumor initiation and progression is crucial.

In this study, the role of WNT signaling in macrophages and its impact on the immunosuppressive TME was investigated using an AOM/DSS-induced inflammatory colon cancer mouse model. *Csf1r.iCre;Porcn^fl/fl^* mice, lacking macrophage-derived WNT, showed increased sensitivity to AOM/DSS-induced colon tumors compared to wild-type (WT) mice. Further analysis revealed a higher number of proliferating cells and an increased presence of Lgr5+ and Dclk+ colon cancer stem cells in the tumors of *Csf1r.iCre;Porcn^fl/fl^* mice compared to WT mice. The elevated expression of WNT target genes in EpCAM-positive sorted tumor cells from *Csf1r.iCre;Porcn^fl/fl^* mice suggests that macrophage-derived WNT does not directly activate WNT signaling in tumor cells. Instead, the absence of WNT release from *Csf1r.iCre;Porcn^fl/^*^fl^ macrophages promotes tumor cell progression by inducing immune evasion, which indirectly leads to the restitution of WNT signaling in tumor cells.

Flow cytometry of tumor cells revealed a higher percentage of PD1 and PDL1 expressing tumor-associated macrophages (TAMs) in *Csf1r.iCre;Porcn^fl/fl^* mice compared to wild-type (WT) mice. Recent reports highlight GSK3β as a potential target for regulating PD1 and PDL1 expression [22, 43]. GSK3β activates the PI3K/AKT signaling pathway by inhibiting PTEN, leading to increased PDL1 expression. Additionally, GSK3β inhibits TBET, a negative regulator of PD1. Canonical WNT signaling, through WNT ligand-receptor interaction, inhibits GSK3β. Our data suggest that the absence of WNT release from TAMs in *Csf1r.iCre;Porcn^fl/fl^* mice abolishes WNT-mediated GSK3β inhibition, which consequently reverses PTEN and TBET activation, leading to upregulation of both PDL1 and PD1 expression. Moreover, the absence of WNT signaling in macrophages promotes an immunosuppressive tumor microenvironment, characterized by the upregulation of PDL1 in T helper cells and myeloid-derived suppressor cells (MDSCs). GeoMx transcriptomics data from tumor tissue further demonstrated that absence of WNT signaling in macrophages induces immunosuppressive gene signature within the tumor microenvironment. These findings indicate that the lack of macrophage-derived WNT in *Csf1r.iCre;Porcn^fl/fl^* mice disrupts the autocrine role of WNT in inhibiting GSK3β within these macrophages. Pharmacological inhibition of GSK3β in *Csf1r.iCre;Porcn^fl/fl^* mice reduced the tumor burden after AOMDSS exposure.

In an ex-vivo co-culture system using Porcupine inhibitor (C59)-treated human macrophages [9], human COLO205 cells, and T cells, a significant upregulation of PDL1 expression in the macrophages was observed. However, this PDL1 upregulation was reversed when C59-treated macrophages were exposed to a GSK3β inhibitor. These observations suggest that WNT autocrine signaling and subsequent inhibition of GSK3β in macrophages are crucial for overcoming their immunosuppressive role. Furthermore, both in vivo and ex vivo experiments suggest that, even with WNT autocrine signaling present, additional pharmacological inhibition of GSK3β in WT macrophages can be beneficial and can be established as therapeutic against colitis induced colon cancer.

Modulating Wnt signaling has indeed been a highly attractive target in colon cancer treatment research for several decades [44, 45]. The Wnt/β-catenin pathway is frequently activated in colorectal cancer (CRC), primarily due to mutations in the APC and β-catenin genes [46, 47]. However, developing successful Wnt-targeted therapies has faced significant challenges and results obtained so far from clinical studies was not promising [48, 49]. Evidence obtained from the present pre-clinical study suggested that global inhibition of WNT/beta catenin signaling may promote immunosuppressive tumor microenvironment as WNT autocrine signaling in macrophages is important to inhibit GSK3beta downstream PD1 and PDL1 expression. Our study also suggested that targeted cell specific pharmacological reprogramming is important compared to global approach. This study therefore established that systemic delivery of macrophage specific liposomal GSK3beta inhibitor can be an effective therapeutic strategy against colitis associated colon cancer.

## Notes

### Competing Interest Statement

The authors have declared no competing interest.

