## Supplementary Figure for "Macrophage-derived WNT regulates tumor immune microenvironment to reduce colitis-associated colon cancer"

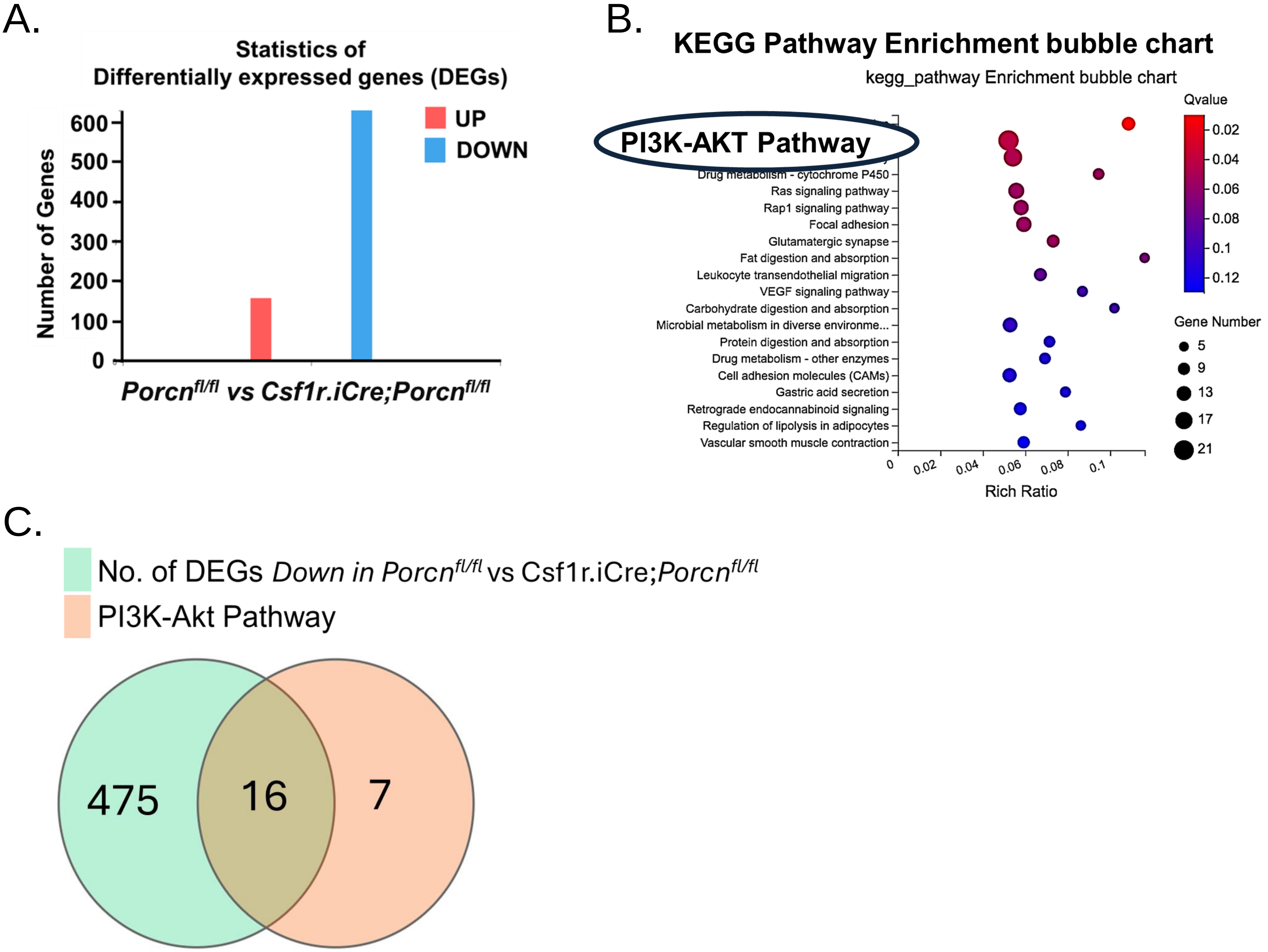


**Supplementary Figure 1. Transcriptomic profiling identifies dysregulated pathways.** A. Statistics of differentially expressed genes (DEGs) showing the number of upregulated (red) and downregulated (blue) genes between *Porcn*$^{fl/fl}$ vs *Csf1r.iCre;Porcn*$^{fl/fl}$ MDSCs. B. KEGG pathway enrichment bubble chart of DEGs. The x-axis indicates the rich ratio, bubble size represents the number of genes. The PI3K-AKT signaling pathway was among the most significantly enriched pathway. C. Venn diagram showing overlap of DEGs between *Porcn*$^{fl/fl}$ vs *Csf1r.iCre;Porcn*$^{fl/fl}$ , with 16 common genes identified, while 7 genes related to Pi3K-Akt pathway.
