## Supplementary Table for "Macrophage-derived WNT regulates tumor immune microenvironment to reduce colitis-associated colon cancer"

**Supplementary Table-1**

**Primer Sequences**

| Gene Name | Forward Sequence (5’-3’) | Reverse Sequence (5’-3’) |
|---|---|---|
| Axin-2 | ATGGAGTCCCTCCTTACCGCAT | GTTCCACAGGCGTCATCTCCTT |
| Ascl2 | TTTCCTGTGCCGCACCAGAACT | CAGCGACTCCAGACGAGGTGG |
| Sox9 | CACACGTCAAGCGACCCATGAA | TCTTCTCGCTCTCGTTCAGCAG |
| Ephb2 | CGCCATCTATGTCTTCCAGGTG | GATGAGTGGCAACTTCTCCTGG |
| Tcf4 | CCTCCAATCCTTCAACTCCTGTG | TCCAAACGGTCTTCGATTCGGC |
| Lef1 | ACTGTCAGGCGACACTTCCATG | GTGCTCCTGTTTGACCTGAGGT |
| Olfm4 | GCCTCCAAAAGTGACCTTGTGC | TGCGTGTGCTGGTGGAAAAGAG |
| Alcam | AGGAACATGGCGGCTTCAACGA | ACACCACAGTCGCGTTCCTACT |
| Lgr5 | AGAGCCTGATACCATCTGCAAAC | TGAAGGTCGTCCACACTGTTGC |
| Bmi-1 | ACTACACGCTAATGGACATTGCC | CTCTCCAGCATTCGTCAGTCCA |
| Epcam | GAGTCCGAAGAACCGACAAGGA | GATGTGAACGCCTCTTGAAGCG |
| CD29 | CTCCAGAAGGTGGCTTTGATGC | GTGAAACCCAGCATCCGTGGAA |
| CD44 | CCAGAAGGAACAGTGGTTTGGC | ACTGTCCTCTGGGCTTGGTGTT |
| CD133 | CTGCGATAGCATCAGACCAAGC | CTTTTGACGAGGCTCTCCAGATC |
| GAPDH | ACCACAGTCCATGCCATCAC | TCCACCACCCTGTTGCTGTA |
